# Genetic diversity of North American popcorn germplasm and the effect of population structure on nicosulfuron response

**DOI:** 10.1101/2023.01.06.523022

**Authors:** Madsen Sullivan, Martin M. Williams, Anthony J. Studer

## Abstract

Popcorn is an important crop in the United States; however, genetic analyses of popcorn are limited and tend to utilize relatively few markers that cannot capture the total genomic variation. To improve the genomic resources in popcorn, a panel of 362 popcorn accessions was evaluated using 417,218 single nucleotide polymorphisms generated using a genotyping-by-sequencing approach. Using this genomic data, a model-based clustering analysis identified two populations. The first comprised North American Yellow Pearl Popcorns and several accessions of the Chilean Curagua landrace. The second, the Pointed and Latin American Popcorns, included all remaining North American (pointed and early popcorns), Latin American, and global accessions. The two populations exhibited large differences in population structure and genetic diversity. The North American Yellow Pearl Popcorns constitute a highly inbred population with limited genetic diversity compared to the Pointed and Latin American Popcorns. Additionally, phenotypic differences between the two populations were observed in kernel color and nicosulfuron sensitivity. A filtered set of SNPs was curated and used for genome-wide association studies and popcorn-specific candidate genes for nicosulfuron tolerance were identified. The genomic characterization described here can be used by breeding programs to accelerate the rate of genetic gain and incorporate genetic diversity into elite popcorn germplasm.

**Core Ideas:**

1. North American Popcorn is composed of two distinct populations that differ genetically and phenotypically.
2. North American Yellow Pearl Popcorns contain limited genetic diversity and are highly inbred.
3. Pointed and Latin American Popcorns exhibit considerable genetic diversity and rapid linkage disequilibrium decay.
4. Kernel color does not affect nicosulfuron sensitivity and instead reflects differences between the populations.
5. Popcorn-specific candidate genes for nicosulfuron tolerance are distinct from dent corn.

## 1 INTRODUCTION

Popcorn (*Zea mays var. everta*) is a popular snack and a valuable commercial crop in the United States. In a global market worth 10 billion USD, the United States has the highest per capita popcorn consumption of any country at over 60 quarts per year, representing a total of 17 billion quarts consumed annually (Serna-Saldivar 2022). Likewise, the United States grows more popcorn than any other nation, with approximately 90,000 hectares of popcorn planted annually, most of which are grown in Nebraska, Indiana, Ohio, and Illinois (USDA NASS, 2022).

Popcorn has been cultivated within the territory of the modern United States since pre-Columbian times, although the range of pre-Columbian popcorn was limited to the American Southwest. Popcorn landraces first reached this region by 4,200 BP (Merrill et al. 2009), likely first from a highland origin, including the landraces Palomero Toluqueño and Palomero de Jalisco, followed by a later mixing with lowland landraces, such as Chapalote and Reventador (da Fonseca et al. 2015; Wang et al. 2017). Although these landraces do pop, the popping quality is poor compared to modern lines in terms of expansion, hull fragmentation, and percentage of unpopped kernels (Vazquez-Carrillo et al. 2019; Bautista-Ramirez et al. 2020).

Despite cultivation in the American Southwest for millennia, popcorn was not widely distributed in North America until the late 18^th^ and early 19^th^ centuries, and even then, the amount grown was limited (Erwin 1950). During the first half of the 19^th^ century, popcorn was typically grown in the United States as a small-scale garden crop for household use (Ziegler 2000). By the late 19^th^ and early 20^th^ century, the popularity of popcorn rose as open-pollinated varieties began reaching more consumers, following the first successful processors and commercial brands in the 1880s and the introduction of the mobile popcorn machine in 1893 (Smith 1999). The predominant types of popcorn grown during this time were the North American Pointed Rice Popcorns or ‘hulless’ types, characterized as having a pointed upper kernel shape and frequently colored white. The first hybrid popcorn used in commercial production was generated using two selections of Japanese Hulless, a white pointed rice variety (Brunson 1937). However, since the mid-20^th^ century, another group of popcorn has largely replaced the pointed rice type commercially – the North American Yellow Pearl Popcorns. This group has a rounded kernel shape and is predominantly yellow, although color variation has been introduced by several breeding programs (Ziegler 2000). While the North American Yellow Pearl Popcorns are the most important for commercial production in the United States, North American Pointed Rice Popcorns are still commercially grown and used in breeding programs.

Characterizing the genetic relationships and identifying variation within these distinct populations of North American popcorn is an important aspect of breeding improved popcorn. Early molecular techniques were used to assess the genetic diversity of North American Yellow Pearl Popcorns and identified three heterotic groups – Supergold, South American, and Amber Pearl (Kantety et al. 1995). Later work by Santacruz-Varela et al. defined three major groups of North American popcorn – the aforementioned North American Pointed Rice Popcorn and North American Yellow Pearl Popcorn, as well as the North American Early Popcorn (2004).

Additionally, each group’s phylogenetic relationship and historical origin were investigated, and possible germplasm sources were identified. The North American Pointed Rice Popcorns appeared to originate from the traditional pointed rice popcorns of Latin America, based on both morphological and genotypic similarities. Furthermore, the North American Yellow Pearl Popcorns were most closely related to, and likely principally derived from, the Chilean Curagua landrace. Finally, the North American Early Popcorns may have been developed from Northern Flint corn and subsequently influenced by European popcorn varieties (Santacruz-Varela et al. 2004). Although this information has been useful in understanding breeding history and relatedness, the limited number of markers and lines has restricted its use in modern breeding programs to heterotic pool development.

Compared with dent corn germplasm, popcorn exhibits considerably inferior agronomic traits, including herbicide tolerance, disease resistance, lodging, yield, and overall vigor (Ziegler 2000). Additionally, differences in herbicide tolerance have been reported within popcorn, with white popcorn generally exhibiting greater herbicide sensitivity than yellow (Loux et al. 2017).

Historically, the poorer agronomic traits of popcorn are due partly to the amount of time and resources breeders have invested relative to dent corn and also because of the emphasis placed on popping quality traits. In addition to agronomic traits, popcorn breeders must consider quality traits directly related to popping and eating quality, including popping expansion, hull separation, frequency of unpopped kernels, texture, and appearance (Matz 1984). Furthermore, dent corn breeding pipelines have integrated molecular characterization and resources, resulting in improved selection and rates of genetic gain, while publicly available resources within popcorn are limited.

High-quality genetic and genomic resources in maize have empowered researchers and facilitated discoveries involving the genetic architecture of traits, candidate gene association and identification, and population structure (Buckler et al. 2009; Cook et al. 2012; Romay et al. 2013; Peiffer et al. 2014; Wallace et al. 2014; Gage et al. 2020). While the genetic resources for mapping populations, diversity panels, and seed banks have been used extensively within dent corn, popcorn germplasm comprises a genetically distinct subpopulation of maize that is underrepresented in these studies. Recently, genotyping-by-sequencing (GBS) was performed on popcorn accessions from CIMMYT (Li et al. 2019; Li et al. 2021) and Chinese breeding programs (Yu et al. 2021) and used to characterize genetic diversity, describe population structure, and perform genome-wide association studies (GWAS). The genomic characterization of these diverse popcorn accessions can be used to assist breeders in incorporating genetic variation into breeding programs and identifying loci controlling important traits in popcorn.

Likewise, the rate of genetic gain in North American popcorn breeding programs would increase dramatically with the incorporation of high-quality genetic and genomic information, as has been utilized in dent corn breeding for decades.

In this study, molecular analysis was performed on 362 publicly available popcorn accessions genotyped with GBS representing the diversity of North American popcorn. The goals of this study were to i) characterize the genetic diversity of North American popcorn, ii) identify population structure within North American popcorn and other groups of popcorn, iii) describe genomic differences between these populations, iv) perform genome-wide association studies, and v) compare findings with previous attempts to characterize North American popcorn.

## 2 MATERIALS AND METHODS

### 2.1 Plant materials and DNA extraction

Of publicly available lines of popcorn, 341 unique accessions were selected from the Germplasm Resources Information Network (GRIN), and 21 accessions were selected from the International Maize and Wheat Improvement Center (CIMMYT). Of these, 79 accessions had available sequence information (Romay et al. 2013). Thus, 283 new popcorn lines were sequenced, with 68 selected for replicate sequencing. Additionally, four dent inbreds with available sequence information were included as outgroup accessions: B73, Mo17, PH207, and Oh43.

Samples were prepared for DNA extraction by bulking tissue from five seedlings of each accession in a 2 mL tube, collected from the youngest leaf of each approximately ten days after germination. DNA was extracted from each sample using a modified CTAB DNA extraction protocol and resuspended in water. DNA was quantified in 96-well plates using PicoGreen (Invitrogen, Carlsbad, CA), and DNA concentrations were normalized to 50ng/μL by diluting with 10 mM Tris, then stored at 4°C.

### 2.2 Genotyping-by-sequencing library construction

Genotyping-by-sequencing libraries were constructed using the protocol described by Elshire *et al*. (2011). Briefly, restriction digestion and ligation were performed in 96-well plates using *Ape*KI by adding 250 ng of DNA from each sample, 1.5μL of 0.1 μM rare barcoded adapter, and 0.5μL of 10 μM common barcoded adapter per well. Libraries were pooled, size-selected and cleaned with Agencourt AMPure XP beads (Beckman Coulter Inc., Brea, CA), and amplified for 15 cycles using KAPA HiFi HotStart ReadyMix (KAPA BIOSYSTEMS). Following a final purification with Agencourt AMPure XP beads, libraries were quantified using PicoGreen. A 1 ng/μL dilution of the library was made using 10 mM Tris, and 5 μL was run on a 1% agarose gel to verify size selection and primer-dimer removal. Fragment size was estimated using a Bioanalyzer 2100 (Agilent, Santa Clara, CA) High Sensitivity DNA Analysis. Pooled libraries were diluted to 10 nmol using 10 mM Tris and sequenced with 100-bp, single-end reads on an Illumina NovaSeq 6000 (Illumina, San Diego, CA). The sequencing data has been submitted to NCBI SRA (PRJNA911304).

### 2.3 Marker calling

The reads from the 79 popcorn accessions previously sequenced by Romay et al. (2013) were downloaded from NCBI SRA (SRP021921) and processed with the new raw sequencing reads using the GBS v2 discovery pipeline from TASSEL 5 (release 5.2.59) (Glaubitz et al. 2014). The pipeline tools and select parameters were run in the following order: First, GBSSeqToTagDBPlugin, with ApeKI and a kmer length of 64, followed by TagExportToFastqPlugin. Next, two alignment indices were built with Bowtie2 (Langmead & Salzberg 2012) using the B73 v5 and HP301 v1 reference genomes (Hufford et al. 2021). Both genome indices and output files were processed using the same parameters afterward. The Bowtie2 alignment command was run using the --very-sensitive preset, resulting in an overall alignment of 90.6% and 91.2% for the B73 v5 and HP301 v1 indices, respectively. The SAM files were then sorted using samtools (Li et al. 2009), and mapping quality (MAPQ) scores were analyzed using Qualimap 2 (Okonechnikov et al. 2016). The SAM files were read using the SAMToGBSdbPlugin with a minimum MAPQ score of 20, after which the DiscoverySNPCallPluginV2 was run using a minimum minor allele frequency of 0.05. Finally, the ProductionSNPCallerPluginV2 was run using the same parameters as the GBSSeqToTagDBPlugin.

### 2.4 Marker coverage and characterization

The genomic distribution of single nucleotide polymorphisms (SNPs) was based on the HP301 v1 reference genome, as the popcorn reads alignments were slightly better than when using the B73 v5 genome. Markers were reported from TASSEL 5 for the Core Analysis set and each chromosome. An additional marker set was generated for imputation and GWAS after filtering out sites with more than 0.75 missing data from the Core Analysis set. Nucleotide substitutions were calculated from the Core Analysis set using the TASSEL 5 GenotypeSummaryPlugin. Markers were identified as intragenic by searching their positions relative to the start and stop positions of the HP301 v1 gene models. The density and distribution of polymorphic SNPs were determined in *R* (R Core Team, 2021) using the *R* package *rMVP* (Yin et al. 2021).

### 2.5 Population structure

A kinship matrix of centered identity by state (IBS) values was calculated for all possible pairwise taxa comparisons in TASSEL. A plot of each pairwise comparison was generated using the *R* package *ggplot2* (Wickham 2016). To formally test for population structure using molecular data, an unsupervised, model-based clustering analysis was performed using ADMIXTURE (Alexander et al. 2009). One through five *K* populations were tested using tenfold cross-validation, with errors minimized at *K* equals two. A second ADMIXTURE analysis was performed in validation after combining the 362 popcorn accessions with 2,284 additional accessions sequenced from the Ames Diversity Panel (Romay et al. 2013). Due to computational restraints, accessions designated stiff stalk or non-stiff stalk were excluded. Reads from the combined set of accessions were aligned to the HP301 v1 reference genome using the TASSEL 5 GBS v2 discovery pipeline without changing any of the aforementioned parameters. In ADMIXTURE, one through twenty *K* populations were tested using fivefold cross-validation, with errors minimized at *K* equals fifteen.

Principal coordinates were generated in TASSEL from the IBS kinship matrix. Three of the coordinates from the matrix with high eigenvalues were visualized using *ggplot2*. After removing 30 accessions with missing data greater than 0.95, phylogenetic analysis was performed using the *R* package *ape* (Paradis et al. 2004) using the neighbor-joining method from a modified Euclidean distance matrix and visualized with the *R* package *ggtree* (Yu et al. 2017). The neighbor-joining algorithm was selected because of the considerable differences in selection and breeding history of the populations included in this study, making alternate methods such as UPGMA untenable. To further investigate the relationship between teosinte and the Pointed and Latin American Popcorn population, an additional neighbor-joining tree was generated. A distance matrix was produced from the seventeen teosinte accessions and ten randomly selected individuals from each of the fifteen ADMIXTURE groups identified from the combined popcorn – Ames Diversity Panel analysis. This distance matrix was used by *ape* to create the second tree and also visualized with *ggtree*.

### 2.6 Population genetics and diversity

To evaluate differences in selection and genetic drift between the two populations, the proportion of monomorphic SNPs within each population was calculated. Both populations were filtered from the Core Analysis set, after which markers in each population were filtered to contain a minimum of two observations. Markers without minor alleles were then filtered out, allowing the number of monomorphic markers to be calculated by subtracting the remaining polymorphic markers. After filtering monomorphic sites, marker minor allele frequencies were calculated from the Core Analysis set using the TASSEL 5 GenotypeSummaryPlugin. The distribution of minor allele frequencies was calculated for each popcorn population.

A marker subset was produced for linkage disequilibrium (LD) analysis by removing SNPs with more than 50% missing data, a MAF less than 0.05, and heterozygosity greater than 0, resulting in 13,031 retained markers. Within each population, LD was calculated between the subset of markers on each chromosome using the TASSEL 5 LinkageDisequilibriumPlugin. The level of LD decay was measured using the squared Pearson correlation coefficient (*r*^2^) between each pair of markers. Pairwise comparisons were binned according to distance using logarithmically increasing bins.

### 2.7 Herbicide treatment

A subset of popcorn accessions was evaluated for response to nicosulfuron at the University of Illinois Crop Sciences Research and Education Center, Urbana, Illinois, during the summer of 2022. A total of 294 genotypes were included in the trial, with 111 North American Yellow Pearl Popcorns and 178 Pointed and Latin American Popcorns. Additionally, five controls were included in the panel – a tolerant sweet corn hybrid (GSS1477), a nicosulfuron-sensitive sweet corn hybrid (Merit), a partially sensitive yellow popcorn hybrid, a tolerant yellow popcorn hybrid, and a tolerant white popcorn hybrid. Genotypes were arranged in a randomized complete block design with three replications. An experimental unit comprised a single 3-meter row containing 10-20 plants.

When the majority of plants reached the V3/V4 growth stage, nicosulfuron was applied to one-half of each row at a rate of 35 g ai ha^-1^ with a 0.25% (v/v) nonionic surfactant and 2.5% (v/v) urea ammonium nitrate. This treatment was delivered using a pressurized CO_2_ backpack sprayer equipped with AI110025 nozzles (TeeJet Technologies, P.O. Box 7900, Wheaton, IL 60187) spaced 51 cm apart on a 1.5-meter boom calibrated to apply 187 L ha^-1^ at 276 kPa. Popcorn plants were evaluated ten days after nicosulfuron application. Injury was assessed visually on a scale of 0-4, with 0 representing no injury and 4 representing plant death.

### 2.8 Imputation and genome-wide association studies (GWAS)

To evaluate the imputation accuracy of different methods, 1% of the GWAS Filtered set was masked and imputed. Imputation was performed with FILLIN (Swarts et al. 2014) and Beagle v5.1 (Browning et al. 2018). FILLIN was run using a haplotype block size of 4,096. Beagle was run using an effective population size of 100,000 and an HP301 genetic map. The genetic map for HP301 was constructed in *R/qtl* (Broman et al. 2003) using GBS data from the HP301 RILs (Elshire et al. 2011), aligned to the HP301 v1 reference genome using the TASSEL 5 GBS v2 discovery pipeline. To compare the two imputation methods, the unmasked, masked, and imputed marker sets were used as input for TASSEL’s ImputationAccuracyPlugin, and error rates were generated.

As the popcorn populations differ in kernel color and response to nicosulfuron, GWAS was performed on both these phenotypes. Both GWAS analyses were performed using a weighted MLM in TASSEL with the FILLIN-imputed marker set, a kinship matrix of normalized IBS values, and a weight matrix of principal coordinates with high eigenvalues. A Bonferroni-corrected p-value of 0.01 resulted in a -log(p) significance threshold of 7.3. Results of both GWAS analyses were visualized using the *R* package *qqman* (Turner 2018).

## 3 RESULTS

### 3.1 Germplasm

To improve the genomic resources and investigate the genetic diversity of popcorn, GBS was performed on 362 popcorn accessions. In GRIN, a total of 640 available accessions were listed as popcorn. Accessions that were also labeled as other kernel types were removed, reducing the count to 379. Then all accessions that appeared to be duplicates based on passport information were removed, bringing the number of unique popcorn accessions down to 304. Of the accessions with multiple kernel types listed, 37 were added back, as they were known popcorn lines. Finally, an additional 21 popcorn landraces from CIMMYT were included, bringing the total number of accessions to 362 (Table S1). Of these, 72 are listed as landraces, 109 as cultivated material, and the remaining 181 as breeding material. In addition to the 362 popcorn accessions, four previously sequenced dent inbreds were included to serve as outgroup lines: B73, Mo17, PH207, and Oh43.

### 3.2 Marker density and distribution

From the GBS data, a Core Analysis set of 417,118 markers with an average minor allele frequency of 0.169 was generated after passing SNP filtering and quality control. A Filtered GWAS set of 200,101 markers with an average minor allele frequency of 0.216 was generated by removing all SNPs with greater than 0.75 missing data in preparation for imputation. Of the nucleotide substitutions from biallelic SNPs, adenine-guanine and thymine-cytosine transitions were the most common (29.5% each), while adenine-thymine transversions were the least common (7.7%). Of the Core Analysis set of markers, 327,876 (78.6%) were identified as intragenic using the HP301 v1 gene models (Hufford et al. 2021), consistent with the observation that SNP density increased from the centromeric to the telomeric regions (Supplementary Fig. S1). Across the 10 maize chromosomes, the Core Analysis set of markers had an average marker density of 197.3 SNPs per Mb, with the highest and lowest marker densities observed on chromosomes 5 (216 SNPs per Mb) and 6 (184 SNPs per Mb), respectively (Table 1). The Filtered GWAS set had an average marker density of 94.6 SNPs per MB, with the highest and lowest densities on chromosomes 1 (108 SNPs per Mb) and 4 (74.7 SNPs per Mb), respectively.

**Table 1.** SNP characteristics summary for the Core Analysis and Filtered GWAS SNP sets

| Chromosome | Chromosome Length (b) | Core Analysis Set |  |  | Filtered GWAS Set |  |  |
| --- | --- | --- | --- | --- | --- | --- | --- |
|  |  | Number of SNPs | SNPs per Mb | MAF | Number of SNPs | SNPs per Mb | MAF |
| 1 | 305,741,490 | 64450 | 210.8 | 0.16807 | 32903 | 107.6 | 0.21527 |
| 2 | 245,520,863 | 51470 | 209.6 | 0.17452 | 24514 | 99.8 | 0.22231 |
| 3 | 240,329,036 | 47791 | 198.9 | 0.17328 | 24554 | 102.2 | 0.22337 |
| 4 | 251,015,817 | 41118 | 163.8 | 0.16409 | 18748 | 74.7 | 0.20173 |
| 5 | 221,904,911 | 47873 | 215.7 | 0.1712 | 23121 | 104.2 | 0.22609 |
| 6 | 177,672,499 | 32691 | 184.0 | 0.16488 | 14358 | 80.8 | 0.21071 |
| 7 | 180,265,189 | 35984 | 199.6 | 0.16534 | 15815 | 87.7 | 0.21834 |
| 8 | 177,329,642 | 34910 | 196.9 | 0.16316 | 16579 | 93.5 | 0.2199 |
| 9 | 164,337,916 | 31699 | 192.9 | 0.1707 | 16053 | 97.7 | 0.21045 |
| 10 | 150,435,773 | 29232 | 194.3 | 0.16794 | 13456 | 89.4 | 0.21866 |
| Total | 2,114,553,136 | 417218 | 197.3 | 0.16872 | 200101 | 94.6 | 0.21576 |

### 3.3 Population structure

To identify relationships among accessions included in the panel, the distribution of IBS pairwise comparisons for the 362 accessions was calculated using the Core Analysis set (Fig. 1). A bimodal distribution was identified, with a large main peak centered on 0.72 and an additional smaller peak centered on 0.82. These results indicate that although most accessions have a weak relationship with other material in this panel, a subset appears to be more closely related.

**Figure 1.**
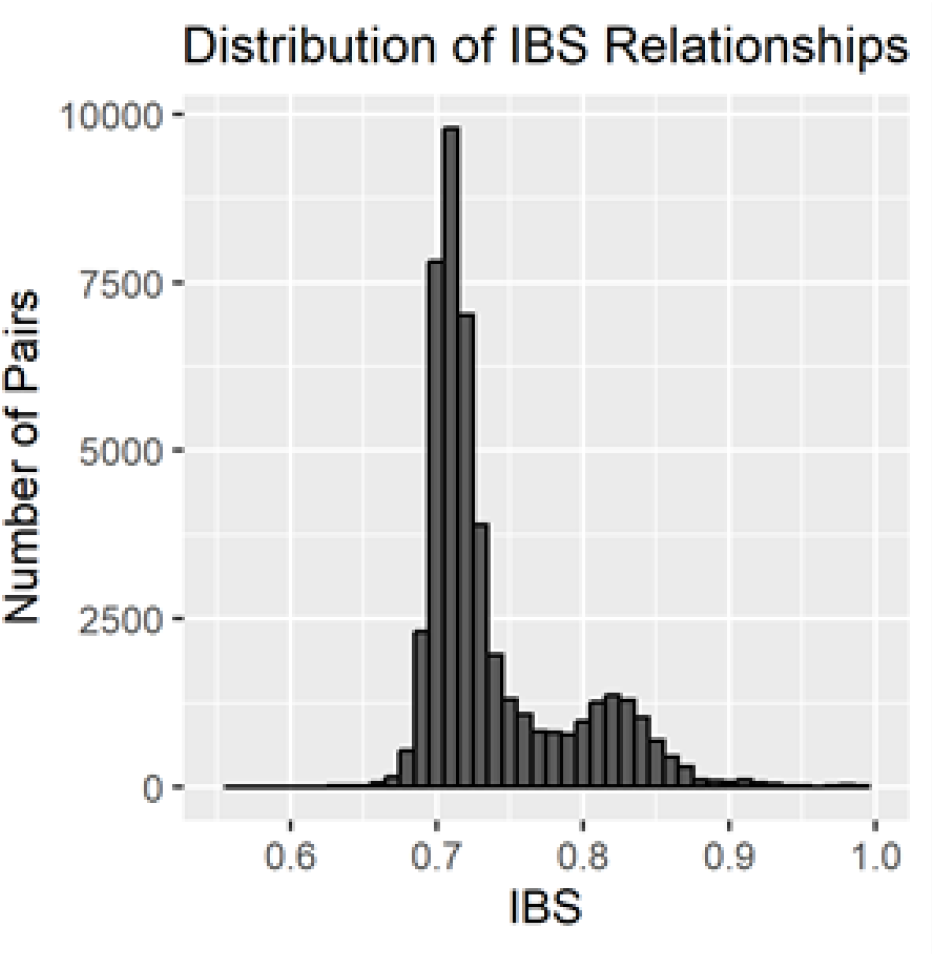
Identity by state (IBS) distribution across 362 popcorn accessions

The North American popcorn populations are comprised of accessions with different breeding histories, geographic distributions, and selection pressures. An ADMIXTURE analysis was performed on all taxa to formally test for population structure using one to five *K* populations.

Cross-validation error was minimized at *K*=2 (Supplementary Fig. S2), and taxa were clustered into two groups of similar size, with 147 individuals assigned to the first group and 215 assigned to the second (Fig. 2, Table S1). Most taxa assigned to the first group are North American Yellow Pearl Popcorns from breeding programs in the United States with known pedigrees and generally fall into one of the previously described heterotic pools. However, several notable taxa from Chile were assigned to this group, including CHZM 05 006, CHZM 06 004, CHZM 06 012, and CHZM 07 097, all of which are members of the Curagua race. The remaining 23 members of Curagua were assigned to the other group of popcorn, which will be referred to as the Pointed and Latin American Popcorns. This group contains many diverse accessions, including all North American Pointed Rice Popcorns (n=21), North American Early Popcorns (n=32), all Latin American pointed and pearl popcorns accessions outside the four members of Curagua (n=98), and nearly all lines developed outside of the Americas (n=64).

**Figure 2.**
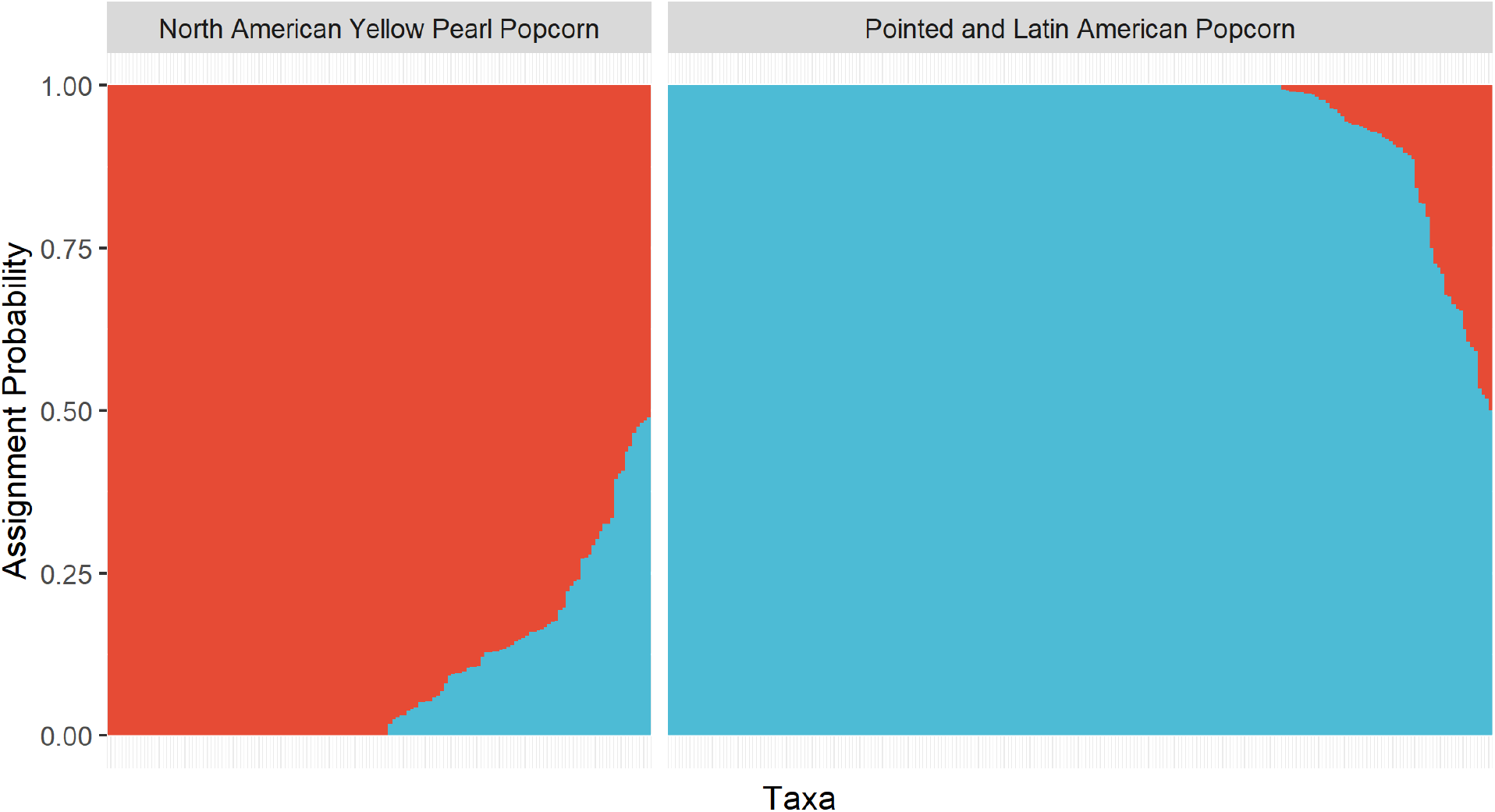
Bar plot of assignment values from ADMIXTURE analysis. Each bar corresponds to one of 362 popcorn accessions and is ordered by the likelihood of assignment. The complete list of assignments can be found in Table S1.

Since more than two popcorn groups have been reported previously in the literature, a second analysis was performed to ensure that the identification of additional popcorn groups was not limited by an insufficient number of related accessions. A combined dataset was generated after merging the 362 popcorn accessions with 2,284 accessions from the Ames Diversity Panel that were not categorized as dent (stiff stalk or non-stiff stalk). After minimizing the ADMIXTURE cross-validation error at *K*=15, the combined 2,646 accessions were assigned to 15 groups, with no additional popcorn groups identified. Although dent lines were excluded, 885 unclassified accessions were assigned to known public subgroups, including B73, B14, B52, Hy, Mo17, W22, Oh43, and WF9, demonstrating that dent material was still well-represented in the analysis. Notably, the analysis assigned the 17 teosinte inbred lines included in the Ames Diversity Panel to the Pointed and Latin American Popcorn group. Thus, the teosinte inbred lines are most genetically similar to the Pointed and Latin American Popcorns, followed by North American Yellow Pearl Popcorns, then sweet corn, and finally dent and flint tropical and temperate material (Supplementary Fig. S3). Other notable taxa assigned to the Pointed and Latin American Popcorn group include accessions of Palomero Toluqueño, Reventador, Chapalote, Pororo, Avati Pichinga, and Pisankalla.

Principal coordinate analysis (PCoA) was also performed to investigate population structure and identify individual taxa grouping, and three of the coordinates explained 83.3% of the variation (Fig. 3). The first coordinate separates North American Yellow Pearl Popcorns from the Pointed and Latin American Popcorns, supporting the *K*=2 optimum number of clusters of the popcorn ADMIXTURE analysis. The second coordinate separates the Pointed and Latin American Popcorns into three general divisions. Most of the North American Pointed Rice Popcorns cluster together near the bottom, with several accessions of Pisankalla. Next, most of the Latin Pointed Rice Popcorns group in the center with members of Curagua and Reventador. Finally, near the top, lowland tropical popcorn landraces such as Avati Pichinga, Pipoca, and Pororo are located together. The third coordinate separates the North American Yellow Pearls, with accessions from the South American heterotic group clustered near the bottom, those from Supergold clustered at the top, and lines developed by Iowa State’s Popcorn Breeding Program located in the center.

**Figure 3.**
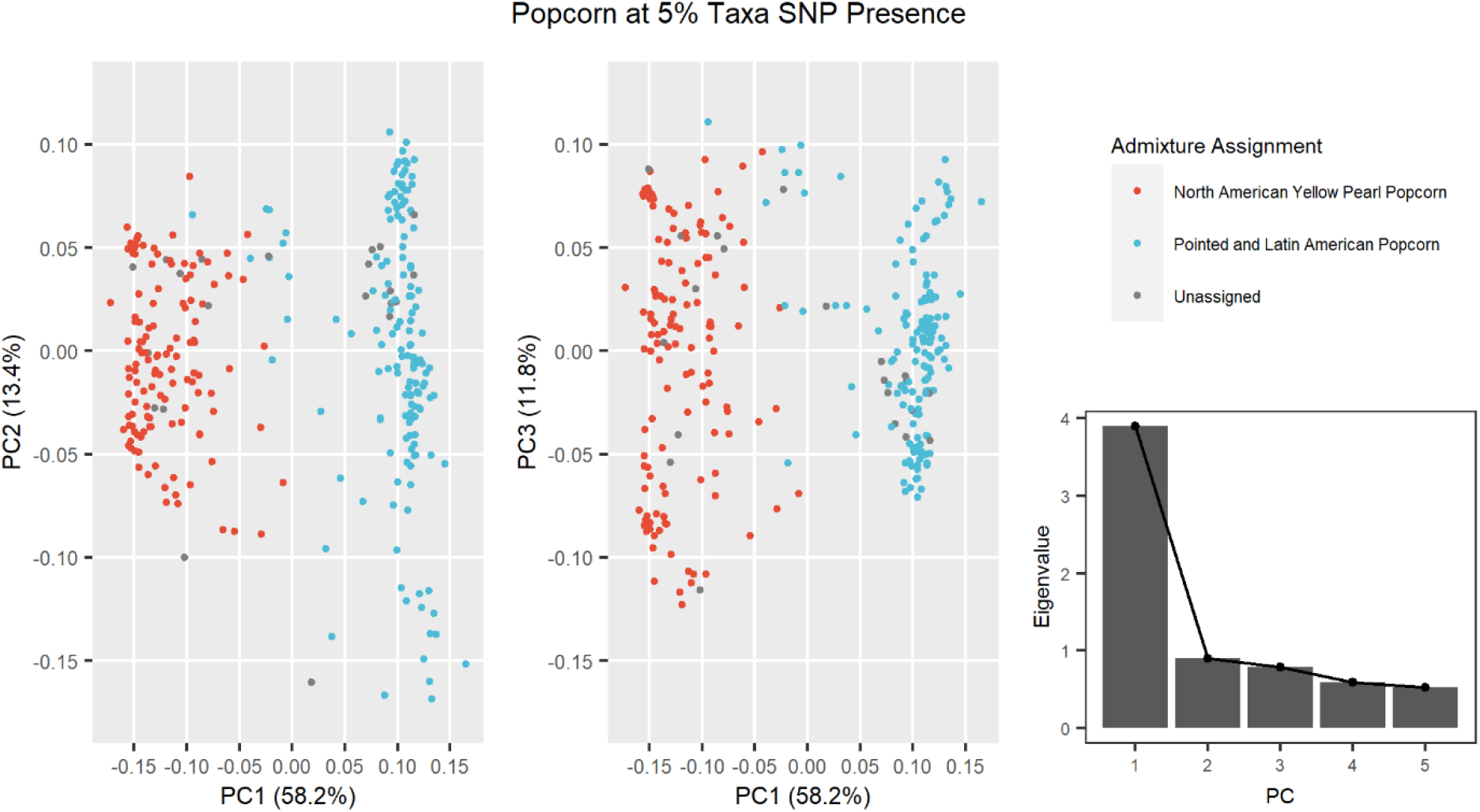
Principal coordinate analysis (PCoA) of 362 popcorn accessions estimated with the Core Analysis set and associated scree plot.

The neighbor-joining tree based on the distance matrix supported the population structure findings of the ADMIXTURE analysis and PCoA (Fig. 4). The major division of clades within the tree corresponded to the separation between North American Yellow Pearl Popcorns and Pointed and Latin American Popcorns. Several Chilean Curagua accessions fall within and basal to the North American Yellow Pearl clade, while the remainder are located within the Pointed and Latin American Popcorns. Two additional Latin American Taxa assigned to the North American Yellow Pearl Popcorn population, BRAZ 2832 and SONO 147, occur within the Supergold clade, indicating that they likely represent a recent movement of North American Yellow Pearl Popcorn germplasm (Fig. 4).

**Figure 4.**
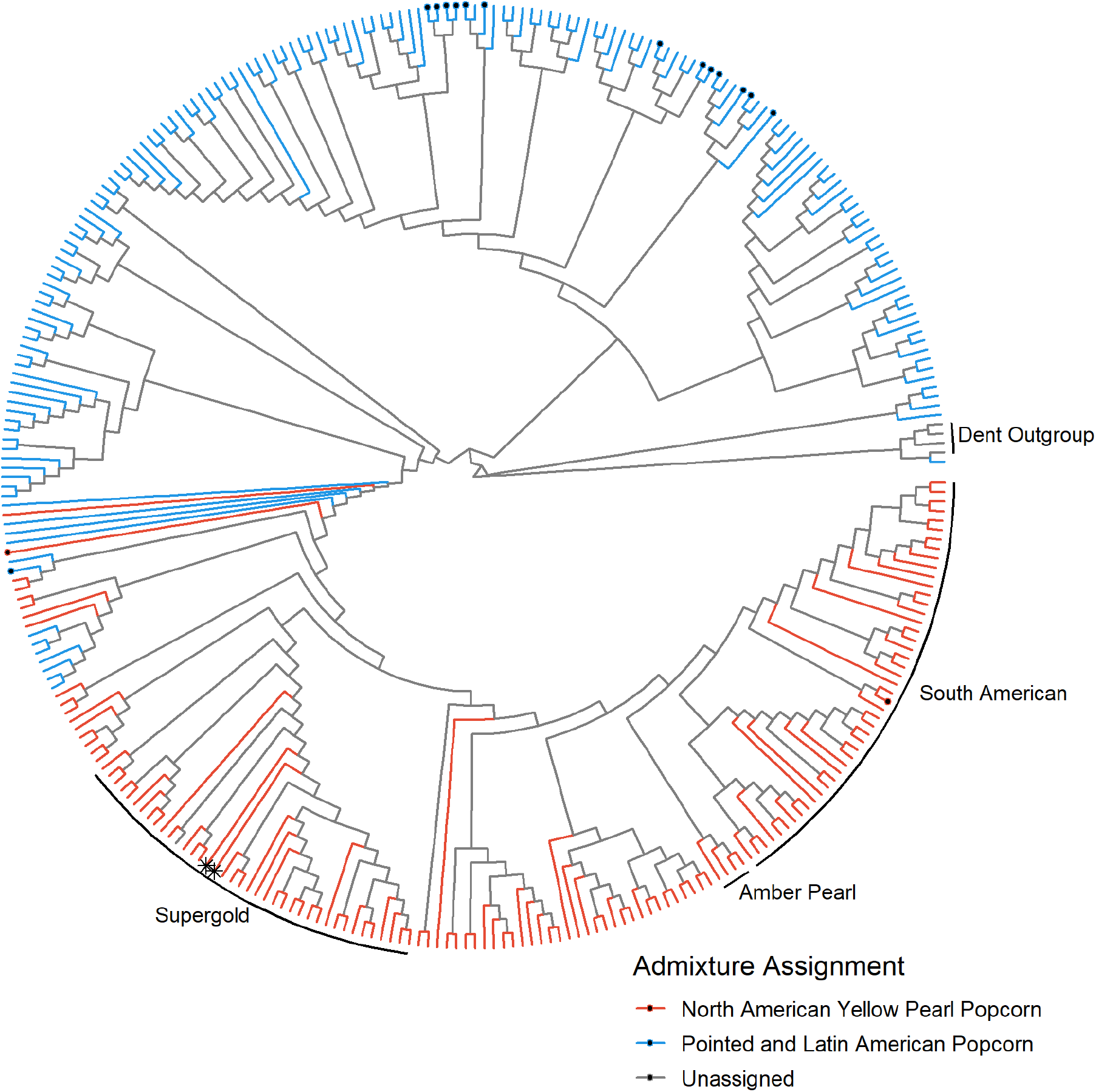
Neighbor-joining tree of 332 popcorn accessions and 4 dent accessions based on a modified Euclidean distance matrix calculated from the Core Analysis set. The three heterotic pools of North American Yellow Pearl Popcorns are labeled, along with the dent outgroup. Accessions SONO 147 and BRAZ 2832 are marked with an asterisk (∗). Accessions belonging to Curagua are marked with a dot (•).

### 3.4 Population genetics and allele diversity

To evaluate differences in allele frequencies and genetic diversity within popcorn germplasm, each population was evaluated for the proportion of monomorphic SNPs, minor allele frequencies (MAF), and LD decay. After filtering markers in each group to account for structural variation, monomorphism was evaluated. The proportion of monomorphic SNPs was considerably different between the two groups, with 5.5 times more monomorphic SNPs in the North American Yellow Pearl Popcorns (0.328) than in the Pointed and Latin American Popcorns (0.059). Additionally, there were differences in MAF between the two populations (Fig. 5).

**Figure 5.**
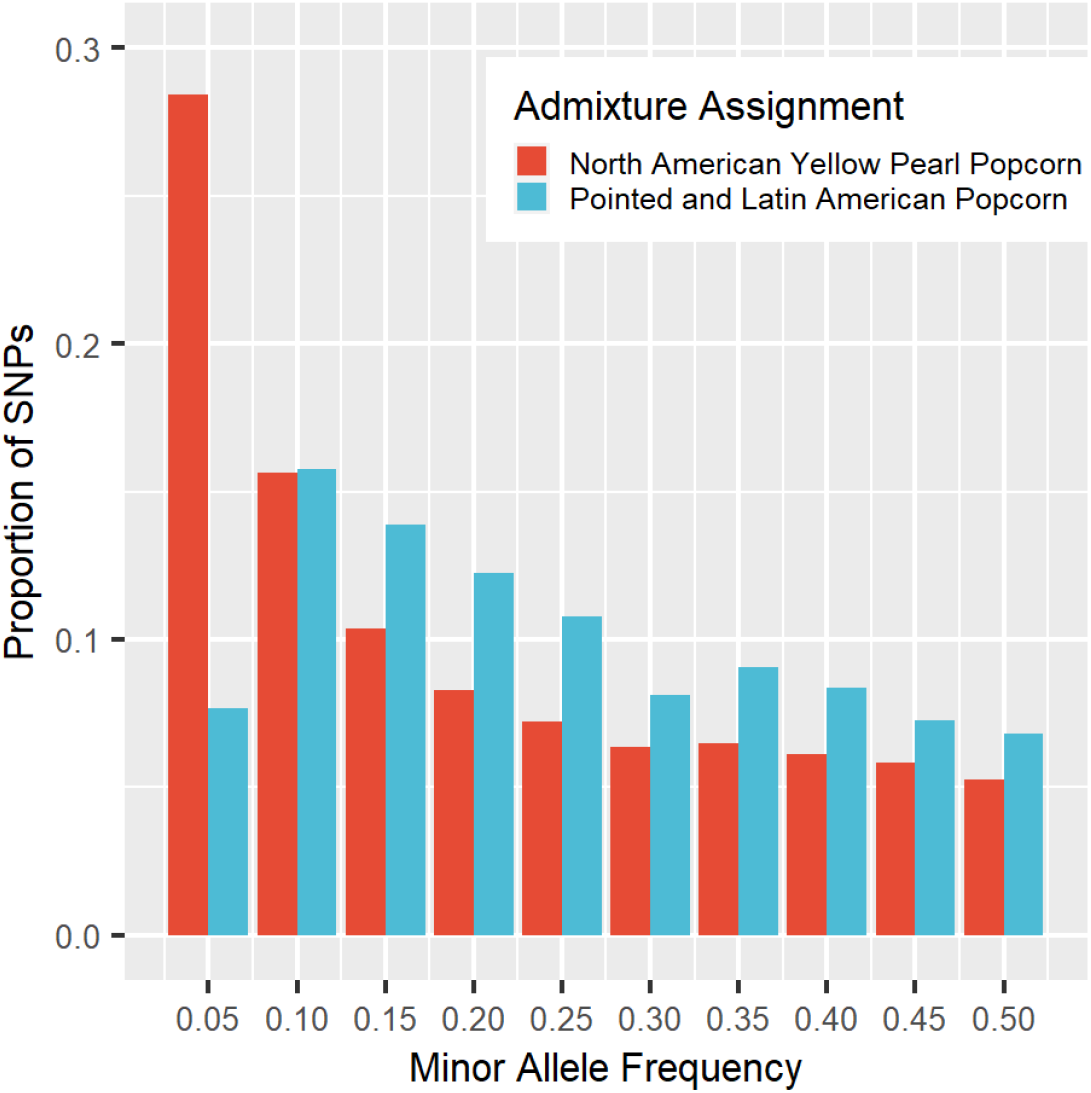
Minor allele frequency (MAF) distributions of the two popcorn populations. Y-axis shows the proportion of SNPs, as differing counts of monomorphic SNPs were removed from each population prior to calculating frequencies.

The relative frequency of SNPs with a MAF of 0.05 or less in North American Yellow Pearl Popcorns and Pointed and Latin American Popcorns were 28.4% and 7.7%, respectively. For all MAF greater than 0.05, the Pointed and Latin American Popcorns contained more SNPs than the North American Yellow Pearl Popcorns. Overall, the North American Yellow Pearl Popcorns contained more SNPs with low MAF than the Pointed and Latin American Popcorns.

Linkage disequilibrium decay also differed between the two populations (Fig. 6). Decay was low in the North American Yellow Pearl Popcorns, reaching a median *r*^2^ of 0.3 within 10 – 99 Kb. In contrast, LD decayed rapidly in the Pointed and Latin American Popcorns, reaching a median *r*^2^ of 0.1 within 10 – 99 bp. When compared with previous evaluations of LD decay in temperate and tropical populations (Romay et al. 2013, White et al. 2020), North American Yellow Pearl Popcorns exhibit slower LD decay than even the B73 heterotic subgroup (*r*^2^ of ∼0.25 within 10 – 99 Kb), while LD decay in the Pointed and Latin American Popcorns was slightly faster than in CIMMYT tropical germplasm (*r*^2^ of 0.1 within 100 – 999 bp).

**Figure 6.**
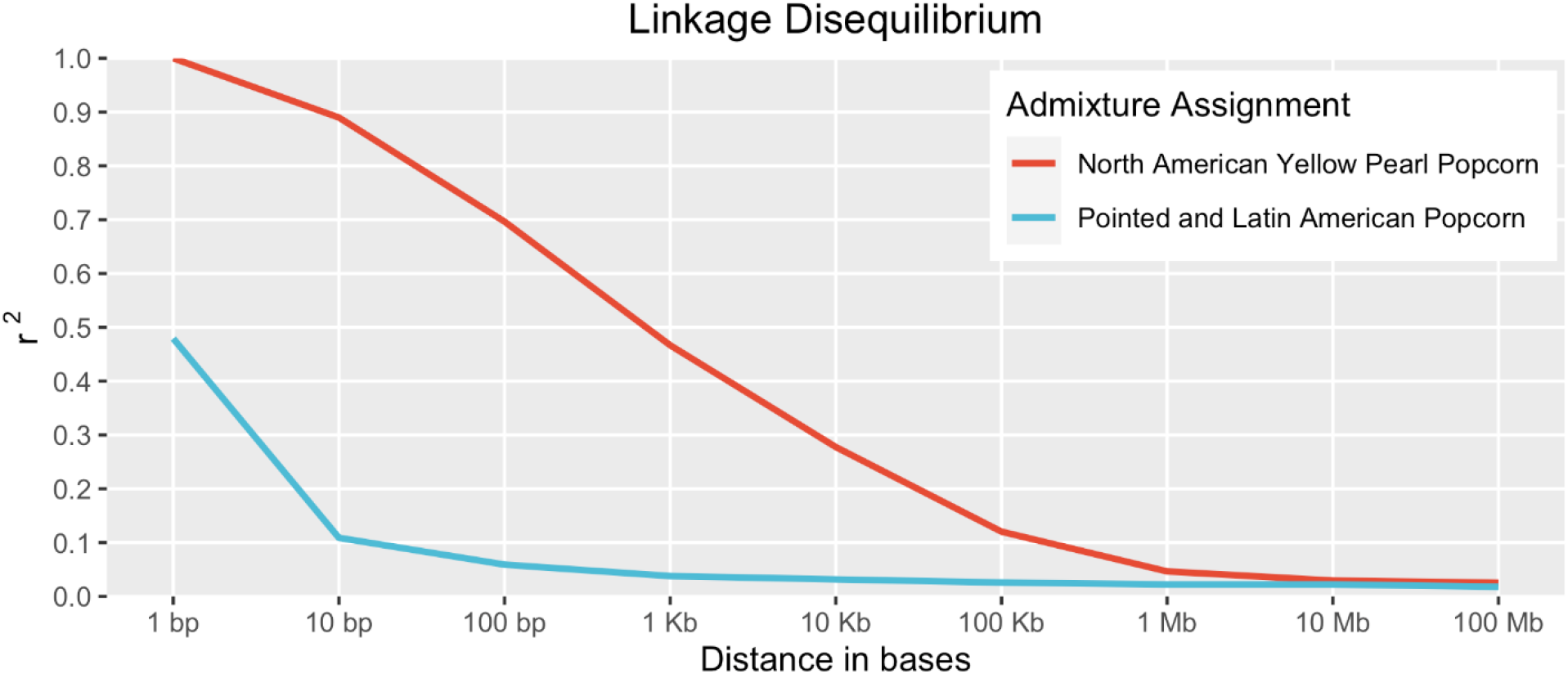
Median linkage disequilibrium (LD) decay of the two popcorn populations. Pairwise comparisons were binned by distance using logarithmically increasing bins.

### 3.5 Genome-wide association studies

A significant limitation of GWAS is the *p*-value correction required to account for multiple tests performed (Tam et al. 2019). Reducing the number of markers tested is one way to ease the stringency of the commonly used Bonferroni correction method. By removing all markers with greater than 0.75 missing data, the resulting Filtered GWAS set reduced the number of markers to approximately half of the Core Analysis set. Furthermore, since the imputation strategies tested here rely on generating haplotypes from existing marker data, any reduction in missing sites improves haplotype generation and subsequent imputation. Two commonly used imputation strategies in maize are Beagle and FILLIN. Both were used to impute the Filtered GWAS set, and error metrics were calculated for each. FILLIN and Beagle had overall error rates of 0.02321 and 0.08173, respectively. Because of the difference in error rates, only the FILLIN-imputed marker set was used to perform GWAS.

The North American Yellow Pearl Popcorns and Pointed and Latin American Popcorns represent two distinct populations with significant differences in population structure, genetic diversity, and breeding history. Additionally, phenotypic differences are known between these two populations, such as kernel color. North American Yellow Pearl Popcorns are typically yellow, with only four white accessions in the panel (3.3%), while the Pointed and Latin American Popcorns have a much higher frequency of white kernels or endosperm, with 111 accessions in the panel (54.7%). Since kernel color is a simple trait that differs among the germplasm assessed here, it was selected as a proof of concept to assess the utility of using this panel for GWAS (Fig. 7). Several significant SNPs were identified, some of which have been associated with kernel color previously (Table S2). The SNP with the strongest association with kernel color is located in *Y1*, which is known to affect the concentration of carotenoid pigments in the endosperm. The SNP with the next strongest association with kernel color is located 200 bases away from Zm00027ab258670, a gene model predicted to have carotene desaturase activity. Three other peak SNPs were identified, none of which have been previously associated with kernel color. These results indicate that despite the known differences in population structure and phenotype frequencies, the panel can identify associations between loci and traits.

**Figure 7.**
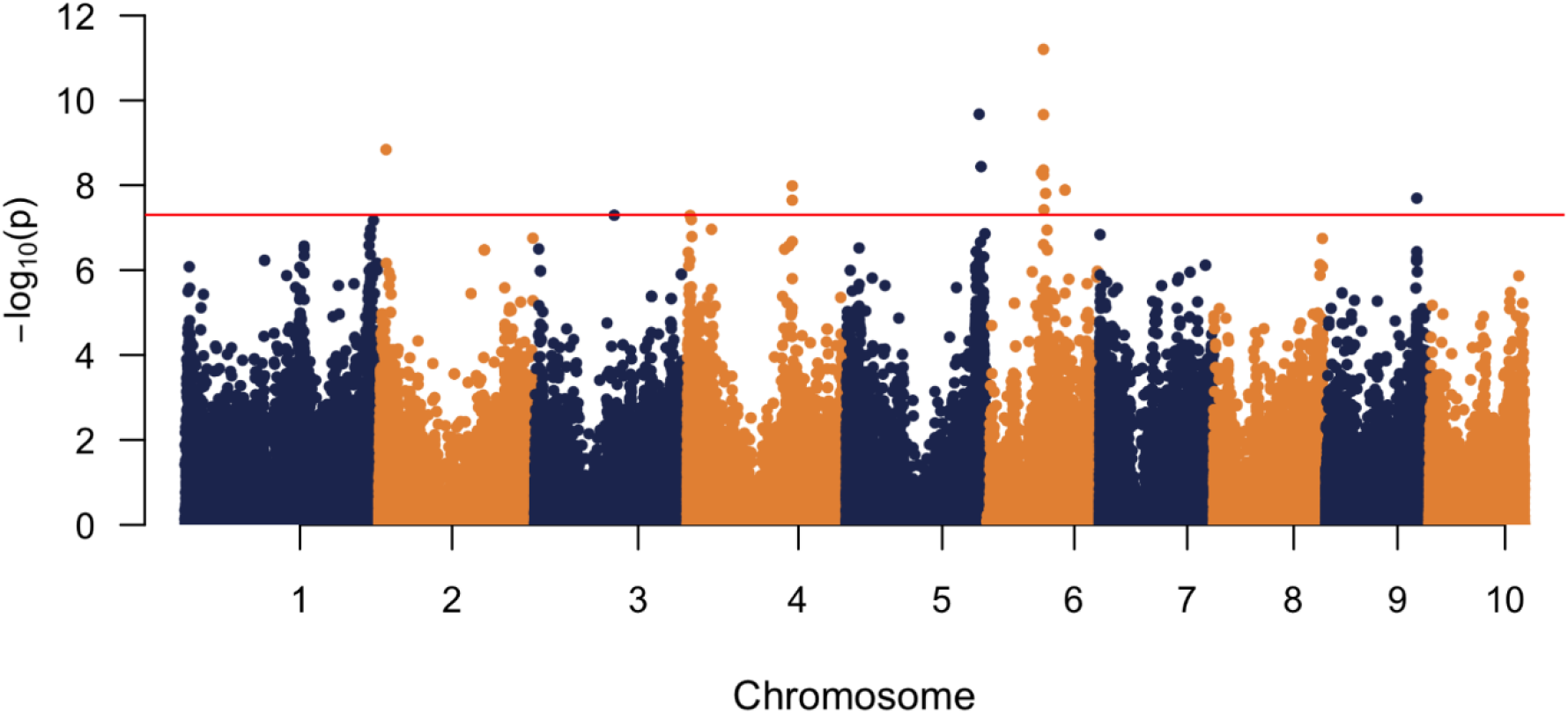
Manhattan plot of genome-wide association study (GWAS) of kernel color from the Filtered GWAS set. The red line represents the threshold after correcting for multiple testing using a Bonferroni correction.

A trait anecdotally associated with kernel color in popcorn is herbicide injury. Herbicide labels often exclude white popcorn from approved use, even when yellow popcorn is included (Loux et al. 2017). While the white color itself should have no bearing on injury, herbicide sensitivity differences between the two popcorn populations may have been reduced to kernel color for simplicity. In the herbicide trial, 4 of the 111 North American Yellow Pearl Popcorns (3.6%) and 32 of 178 Pointed and Latin American Popcorns (18%) showed typical nicosulfuron injury (Table S3). Of the 32 injured Pointed and Latin American Popcorns, several are important historical lines grown in the American Midwest, including Japanese Hulless, Australian Hulless, H5505, IA DS 38-W, and IA DS 43-W. Notably, these pointed lines are all white and represent much of the variation of the pointed rice popcorns traditionally grown in North America.

The Filtered GWAS set was used to map nicosulfuron injury. A total of nine SNPs representing seven peaks were found to be significant (Fig. 8, Table S4). The SNP with the strongest association with injury is located in *cl11433_2b*, a transcription factor associated with the thylakoid. The next strongest association was with a subunit of NADH dehydrogenase. Other SNPs with statistically significant associations include a predicted protein kinase, the vacuole-associated phospholipase *pco139881*, a locus 1.3 kb upstream from a predicted aminoacyl-tRNA synthetase, the cytochrome p450 *CYP72A27*, and a predicted monooxygenase glycosyltransferase. Although these genes have not previously been implicated in nicosulfuron injury, many of the classes to which they belong are associated with herbicide tolerance.

**Figure 8.**
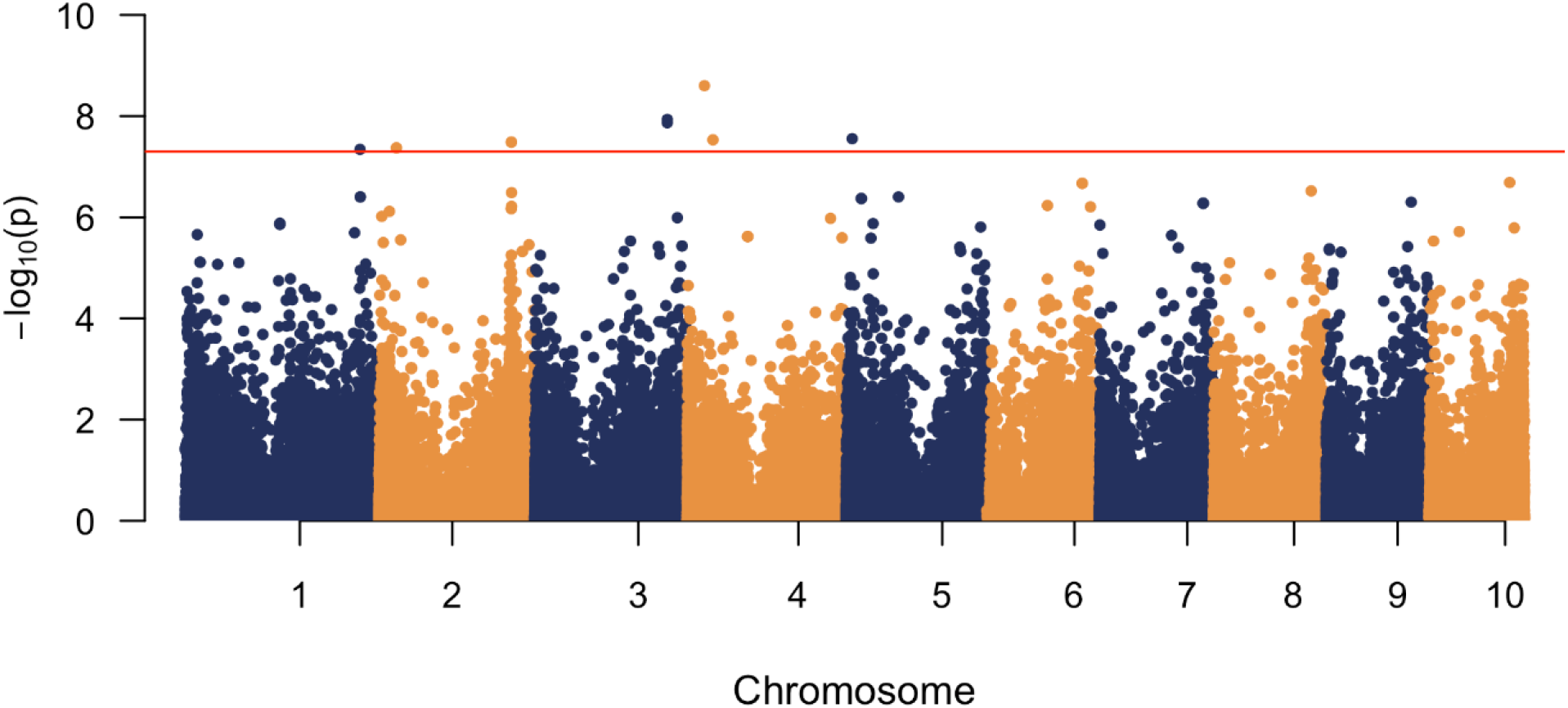
Manhattan plot of genome-wide association study (GWAS) of nicosulfuron injury from the Filtered GWAS set. The red line represents the threshold after correcting for multiple testing using a Bonferroni correction.

## 4 DISCUSSION

### 4.1 Population Structure

The results of IBS, ADMIXTURE, PCoA, and the neighbor-joining tree support two populations of popcorn in the panel investigated here: the North American Yellow Pearl Popcorns and the Pointed and Latin American Popcorns. Both the PCoA and neighbor-joining tree indicate a degree of sub-grouping within each group, but these subgroups were not significantly different based on the genomic data. Within the North American Yellow Pearl Popcorns, differences in population structure of the Supergold and South American heterotic groups were apparent, and there was a minor grouping effect on lines originating from Iowa State’s Popcorn Breeding Program. Divisions within the Pointed and Latin American Popcorns resulted in three general subgroups – North American Pointed Rice Popcorns, Latin Pointed Rice and highland popcorn landraces, and lowland tropical popcorn landraces. It is important to note that not all individuals typically assigned to one of these subgroups clustered with their subgroup, though the majority do. Although sub-groups were identified in both populations, they are not comparable in magnitude to the division between North American Yellow Pearl Popcorns and Pointed and Latin American Popcorns. Therefore, two groups are proposed here that explain most of the variation in population structure in this popcorn diversity panel.

Foundational work on the phylogenetic relationships of North American popcorn identified more than two groups of popcorn. After evaluating 56 popcorn populations for morphological traits, isozymes, and SSRs, six major divisions were previously identified (Santacruz-Varela et al. 2004). Two groups of pointed rice popcorn of North American and Latin American origin were proposed, followed by the related North American Early Popcorns. A single population of North American Yellow Pearl Popcorns was identified, with minimal divisions between the heterotic groups. Finally, two additional groups of Latin America were identified, one corresponding to lowland races of Argentina, Brazil, and Paraguay, and the other composed of various accessions from Chile, Uruguay, and Mexico.

Besides the number of lines, a major difference between previous studies and the results presented here is that morphological traits were not included when determining phylogenetic relationships and population structure. All traits are composed of genotypic and environmental effects, and many of the vegetative and phenological traits included in previous analyses of popcorn are known to be affected by photoperiod sensitivity, including plant height, ear height, leaf number, and days to flowering (Fei et al. 2022). As few as four genomic regions have been identified that control photoperiod responses in these traits (Coles et al. 2010). Therefore, including these traits in measures of phylogenetic divergence likely does not reflect genomic differences between populations and could be heavily influenced by photoperiod sensitivity. This is supported by the conclusion that the six previously identified popcorn populations were primarily assigned based on results of the morphological characterization (Santacruz-Varela et al. 2004). The inclusion of morphological traits that do not reflect genome-wide differences explains the discrepancy between the six populations of popcorn identified previously and the two populations presented here.

Investigation into the genetic diversity and population structure of CIMMYT accessions and Chinese germplasm has identified distinct popcorn populations in their panels. A panel composed of teosinte, popcorn landraces, non-popcorn landraces, North American Yellow Pearl Popcorns, and elite CIMMYT inbreds was used to identify loci associated with temperate and tropical adaptation (Li et al. 2019). This panel identified two groups of popcorn, one corresponding to the North American Yellow Pearl Popcorns and the other associated with landraces from Latin America. The genetic diversity and population structure within a popcorn collection of Chinese accessions, CIMMYT landraces, and inbreds from the United States were recently characterized (Yu et al. 2021). In this study, popcorn accessions were assigned to one of three groups – Latin American landraces, North American Yellow Pearl Popcorns, and recently developed inbreds from breeding programs, likely of North American Yellow Pearl origin. Based on these studies and the findings presented here, North American Yellow Pearl Popcorns have consistently been shown to constitute a genetically distinct pool of germplasm compared to Latin American landraces.

In contrast with the uniform separation of North American Yellow Pearl Popcorns from other types of maize, the assignment of the Pointed and Latin American Popcorns is not as consistent. Single-group assignment of popcorn landraces of Latin America has been previously reported when compared with Chinese germplasm as discussed above (Yu et al. 2021), Brazilian popcorn inbreds (Vittorazzi et al. 2018), and highland and lowland Argentine landraces (Bracco et al. 2012). However, an investigation of Latin American and Caribbean landraces identified sixteen populations, three of which included popcorn (Bedoya et al. 2017). The first of the three populations contained Palomero Toluqueño, Chapalote, and Reventador, all of which are considered primitive popcorns of Northern Mexico. The second group was related to the first and contained Canguil, Confite Puneño, and Pisankalla, all of which are pointed. The third group was unrelated to the first two and contained lowland Avati popcorn. More recently, an analysis of 575 popcorn landraces from Latin America identified nine genetic clusters using discriminant analysis of principal components (DAPC) without considering a priori groups (De Almeida Silva et al. 2020). However, it is important to consider differences in methods when comparing these findings.

Most popcorn groups have been identified using model-based clustering approaches such as STRUCTURE (Pritchard et al. 2000) or ADMIXTURE. Furthermore, model-based clustering methods assign individuals to populations relative to the entire panel – populations identified by an initial clustering analysis can be further divided into sub-groups by subsequent clustering analyses, as with the American and Caribbean landraces discussed above. Additionally, methods based on PCA, like DAPC, tend to provide higher estimates of the total number of populations than model-based clustering approaches (Intarapanich et al. 2009, Linck & Battey 2019) and are especially sensitive to the influence of a priori vs. de novo groupings (Miller et al. 2020).

Because of these considerations, a single assignment by the model-based clustering method ADMIXTURE on the Core Analysis set was used to assess population structure for the entire panel. The findings reported here largely agree with similar approaches to estimate population structure in popcorn, and differences with other reports largely stem from assignment methods.

### 4.2 Genetic diversity

The Pointed and Latin American Popcorns contain a variety of accessions, including pointed popcorns from the American Midwest. It is unlikely that the genetic similarities between lines of pointed popcorn from North America and Latin America indicate a recent introduction or transfer of germplasm between continents, as the lines constituting this group were geographically widely distributed. Instead, these similarities may indicate a pre-Columbian interchange of popcorn between North and South America, as has been previously suggested (Bedoya et al. 2017). Additionally, the genetic similarity may reflect reduced admixture with other landraces due to the high frequency of the dent-sterile *Ga1-S* allele in North and South American popcorn (Goodman et al. 2021). This aligns with the finding that popcorn landraces tend to group separately from other maize types, as discussed above.

North American Yellow Pearl Popcorns share genetic similarities with several Curagua accessions from Chile. Genetic similarity between United States varieties and South American material has been noted before (Vigouroux et al. 2008, van Heerwaarden et al. 2010). However, this has generally been attributed to the post-Columbian movement of germplasm from the United States to South America. Many Chilean, Argentine, and Brazilian races are of United States origin, including Dulce Golden Bantam, Hickory King, and Dente Branco. The presence of Curagua accessions in the North American Yellow Pearls is evidence that intercontinental movement from South to North also occurred.

The North American Yellow Pearl Popcorns have a much higher LD and proportion of monomorphic markers than Pointed and Latin American Popcorns and other populations of maize (Romay 2013). North American Yellow Pearl Popcorns also exhibit higher IBS values when compared with other North American Yellow Pearl Popcorns and lower minor allele frequencies, indicating a high rate of inbreeding and lower genetic variation. Considering the few Curagua accessions clustered among the North American Yellow Pearls (Fig. 4), this reduced genome variation is consistent with a small founder population and fewer meiotic events relative to the considerably older Pointed and Latin American Popcorns. The genetic variation within the three accessions of Curagua assigned to the North American Yellow Pearls represents only a portion of the total variation present in all members of the Curagua race. If members of Curagua were transported to New England by sailors, as previously suggested (Smith 1999), they would have likely contained a limited gene pool, and subsequent inbreeding beginning in the 1940s would have exacerbated the limited genetic diversity. However, it appears that after development in the United States, some of this germplasm returned to Latin America, as evidenced by the SONO 147 and BRAZ 2832 accessions (Fig. 4). Although little additional information is known about BRAZ 2832, the passport data for SONO 147 (National Genetic Resources Program, 2022) contains the alternate name “Reventador Gringo”, suggesting that growers associated this population with non-native material.

### 4.3 Kernel color, popcorn populations, and nicosulfuron injury

The product of the *Y1* gene catalyzes the first step in the maize carotenoid biosynthesis pathway. A SNP in this gene has been identified as a causal mutation that results in a Thr to Asn substitution (Fu et al. 2013). The peak SNP identified in the GWAS is tightly linked with the causal SNP, separated by 346 bases. Another SNP occurs close to a gene model predicted to have carotene desaturase activity and is located 3 Mb away from a locus previously associated with popcorn flake color in CIMMYT germplasm (Li et al. 2021). Given the structural variation between popcorn germplasm and most material used to evaluate kernel color, it is plausible that additional loci may be identified outside the core genome (Lu et al. 2015, Hufford et al. 2021). The three remaining SNPs reported here have not been previously associated with kernel color, although two occur in known genes, *wrky11* and *nc3*, and the other in an uncharacterized gene. Further investigation is necessary to determine whether these additional associations represent biologically meaningful relationships or spurious correlations from population structure.

Recent studies have been unable to identify differences in herbicide sensitivity among several (i.e., up to eight entries) commercial white and yellow popcorn hybrids (Barnes et al. 2020a; Barnes et al. 2020b). However, given the unknown origin of the hybrid parents used in these studies and the prevalence of North American Yellow Pearl Popcorns in commercial germplasm, these hybrids were likely selected using kernel color alone and do not represent the two North American popcorn populations. The results presented here show significant differences in the two popcorn populations for susceptibility to nicosulfuron. Kernel color has no bearing on herbicide injury and demonstrates the importance of population structure in genetic analyses. Kernel color represents a major phenotypic difference between the generally nicosulfuron-tolerant North American Yellow Pearl Popcorns and the more susceptible Pointed and Latin American Popcorns.

Nicosulfuron, along with other sulfonylurea herbicides, inhibits acetolactate synthase (ALS), the first enzyme in the biosynthesis of branched-chain amino acids (BCAAs). Although encoded by the nuclear genome, ALS is localized in the chloroplast (Mazur et al. 1987). Along with other enzymes involved in BCAA biosynthesis, ALS interacts with many substrates and products of cellular respiration, including pyruvate, NAD, and FAD (Duggleby et al. 2008). While target-site resistance of sulfonylurea herbicides has been demonstrated in maize (Anderson & Georgeson 1989; Li et al. 2020), non-target site resistance is more common and complex (Délye 2013).

Non-target site resistance involves initial detoxification, often in the form of cytochrome p450s and other oxidases, followed by conjugation of thiols or glucosyl groups, and subsequent transport and degradation in the vacuole or extracellular spaces (Yuan et al. 2007).

In sweet and dent corn, detoxification of several post-emergent herbicides, including nicosulfuron, is driven largely by *CYP81A9*, a cytochrome p450 produced by the *Nsf1* locus (Nordby et al. 2008, Choe and Williams 2020). Although this does not appear to be the case with nicosulfuron response in popcorn, other candidate genes were identified. Since the pool of available NADH in oxidative phosphorylation is driven largely by pyruvate, variation in NADH dehydrogenase may allow plants to tolerate changes in pyruvate as ALS becomes bound by nicosulfuron. Additionally, variation in glycosylation and changes in the vacuole is consistent with non-target site resistance. Furthermore, altered amino acid synthesis and phosphorylation may be involved in the alternate metabolic flux of some BCAAs (Joshi et al. 2010). Finally, as many of these processes occur within, or are related to, the chloroplast, changes in the transcription factors of chloroplast-associated genes would be expected.

Although *CYP72A27* is a cytochrome p450, its identification here is somewhat unexpected, as it belongs to the CYP clan 72 group of cytochrome p450s. Members of this clan are generally associated with secondary metabolism, while clan 71 enzymes, such as *CYP81A9*, are responsible for herbicide detoxification (Prall et al. 2016; Brazier-Hicks et al. 2022). However, sulfonylurea herbicide metabolism via clan 72 enzymes has been shown in other crops, such as *CYP72A31* in *Oryza indica* (Saika et al. 2014) and *CYP749A16* in *Gossypium hirsutum* (Thyssen et al. 2018). Furthermore, several members of CYP72 were shown to be differentially expressed in nicosulfuron tolerant and susceptible lines of maize when treated with the herbicide (Liu et al. 2015). Alternatively, the role of *CYP72A27* in detoxifying nicosulfuron may be unique to the Pointed and Latin American Popcorns and simply not previously reported. However, it is possible that rather than directly detoxifying nicosulfuron, *CYP72A27* and other genes identified here respond more like safener-induced enzymes, conferring protection via a stress signaling pathway (Riechers et al. 2010). Finally, as with other candidate genes identified using GWAS, *CYP72A27* may have a spurious correlation with nicosulfuron response. Determining the roles and mechanisms of the genes presented here with nicosulfuron tolerance would require further investigation, including the phenotyping and genotyping of additional members of the Pointed and Latin American Popcorns.

## 5 CONCLUSION

High-quality genetic resources are important tools for maize breeding programs. A collection of 362 popcorn accessions were analyzed here using 417,218 SNPs. Two distinct populations of popcorn were identified that exhibit genetic and phenotypic differences. North American Yellow Pearl Popcorns constitute a pool of germplasm with reduced genetic diversity and greater rates of inbreeding when compared with the Pointed and Latin American Popcorns. Kernel color frequencies were shown to differ between the two populations, along with sensitivity to nicosulfuron. Novel candidate genes responsible for nicosulfuron tolerance were identified in popcorn. The genomic characterization provided here should be incorporated into popcorn breeding programs to direct improvement and accelerate the rate of genetic gain. Additionally, this collection of popcorn germplasm and genetic data can be used by the maize research community to explore the genetic architecture of traits in popcorn and be compared with loci identified in other populations of maize.

## Abbreviations

ALS: acetolactate synthase
BCAA: branched-chain amino acid
CIMMYT: International Maize and Wheat Improvement Center
DAPC: discriminant analysis of principal coordinates
GBS: genotyping-by-sequencing
GRIN: Germplasm Resources Information Network
GWAS: genome-wide association studies
IBS: identity by state
LD: linkage disequilibrium
MAF: minor allele frequency
MAPQ: mapping quality
PCoA: principal coordinate analysis
SNP: single nucleotide polymorphism

## SUPPLEMENTAL MATERIAL

Supplemental Table S1 ADMIXTURE assignment probabilities of 362 popcorn accessions by population.

Supplemental Table S2 Significant markers identified by GWAS of kernel color.

Supplemental Table S3 Nicosulfuron injury of 294 accessions, including 111 North American Yellow Pearl Popcorns, 178 Pointed and Latin American Popcorns, and five controls.

Supplemental Table S4 Significant markers identified by GWAS of nicosulfuron injury.

Supplemental Figure S1 SNP density and distribution from the Core Analysis set across chromosomes at a 1Mb window size.

Supplemental Figure S2 Admixture cross-validation plot for 362 popcorn accessions using the Core Analysis set.

Supplemental Figure S3 Neighbor-joining tree of 150 maize and 17 teosinte accessions based on a modified Euclidean distance matrix calculated from the Core Analysis set.

## ACKNOWLEDGMENTS

The authors thank Nicholas Hausman for managing the field design, planting, and nicosulfuron application, Crookham Company for the popcorn hybrids used in the herbicide trial, and Marlee Labroo, Lucas Roberts, and Lynn Doran for assistance with library preparation.

## AUTHOR CONTRIBUTIONS

Madsen Sullivan: Conceptualization, Data curation, Formal analysis, Investigation, Methodology, Software, Validation, Visualization, Writing – original draft, Writing – review & editing. Martin M. Williams II: Conceptualization, Methodology, Resources, Writing – review & editing. Anthony J. Studer: Conceptualization, Funding acquisition, Methodology, Project administration, Resources, Supervision, Writing – original draft, Writing – review & editing.

## CONFLICT OF INTEREST

The authors declare no conflict of interest.

## Notes

### Competing Interest Statement

The authors have declared no competing interest.

### Summary of Updates

Accession IDs updated

